# Historical plant embryos as alternative sources of ancient DNA for whole genome sequencing

**DOI:** 10.64898/2026.02.25.707975

**Authors:** Hong Phuong Le, Stefano Porrelli, Yoon Kyung Lee, Simon Juraver, Flora Pennec, Mark Nesbitt, Koji Numaguchi, Rafal Marek Gutaker

**Affiliations:** Royal Botanic Gardens, Kew, Richmond, Surrey, United Kingdom; Department of Agriculture, Forestry and Bioresources, Plant Genomics and Breeding Institute, Research Institute for Agriculture and Life Sciences, Seoul National University, Seoul, South Korea; Direction Générale Déléguée aux Collections, Muséum Nationale d’Histoire Naturelle, Paris, France; UMR 7206 Eco-anthropologie (CNRS-MNHN-Université Paris Cité), Paris, France; Graduate School of Agricultural Science, Kobe University, Rokkodai 1-1, Kobe, Japan

## Abstract

Natural history and agricultural collections, which contain hundreds of millions of specimens classified in terms of time, space, and taxonomy, are valuable resources for diverse fields of research. Since the first success of ancient DNA (aDNA) isolation in the 1980s, these repositories, including herbaria for plants, have been intensively used to support studies in taxonomy, macroevolution, and genetic responses to anthropogenic activities over the past centuries. Two major challenges of aDNA research are environmental contamination and DNA degradation. For herbarium specimens, aDNA is usually extracted from leaf samples. It is highly fragmented (typically length of 50 to 100 bp) with a higher breakdown rate than that in most bone remains. To optimise the amount of data retrieved and minimise destructive sampling, we isolated DNA from an unconventional plant tissue type – seed embryos. We carried out whole-genome sequencing and compared sequenced DNA quality between embryo and leaf tissue. We evaluated endogenous DNA proportion, median fragment length, damage fraction per site (*λ*), decay rates, nucleotide misincorporations, and library complexity for three species: cultivated rice *Oryza sativa*, wild rice *O. rufipogon*, and wild barley *Hordeum spontaneum*. In *O. sativa*, embryos exhibited significantly higher endogenous content and median fragment length than leaves, while in *O. rufipogon* only median fragment length was higher. The superior DNA preservation was likely due to the protective role of the seed husk, which might play an important role in DNA preservation in plants collected in the tropics. By contrast, in temperate *H. spontaneum*, tissue type had minimal impact on DNA quality. Despite the minuscule size of the embryos, all derived genomic libraries were highly complex, sufficient for deep whole genome sequencing. These results highlight seed embryos as a promising alternative aDNA source for millions of herbarium specimens, and enable effective genomic analyses of other historical plant collections, such as economic botany and anthropological museum collections.

## INTRODUCTION

Herbarium specimens—pressed, dried plants mounted on paper—represent one of the largest and most comprehensive biological archives available for research. Globally, nearly 4,800 herbaria house over 400 million specimens, encompassing all known plant taxa from all continents (*Index Herbariorum - The William & Lynda Steere Herbarium*, n.d.). These collections are invaluable for a wide range of research areas, including taxonomy, biodiversity, and morphology (Heberling et al., 2019). With advances in molecular biology and next-generation sequencing (NGS), herbaria, along with other historical plant collections, have become increasingly important as genetic resource repositories, supporting studies of plant evolution, phylogeny, and ecological interaction (Burbano & Gutaker, 2023; Kistler et al., 2020).

However, DNA isolated from herbarium specimens is reported to be highly fragmented and chemically damaged (Gutaker & Burbano, 2017). Following plant collection and cell death, DNA is initially degraded by intracellular nucleases and microbial activities. Over time, hydrolytic and oxidative reactions further reduce DNA integrity at a slower pace (Lindahl, 1993). DNA degradation and microbe colonisation occur during both the preparation process of herbarium specimens and storage. Post-collection treatment of specimens usually involves heat for desiccation, freezing temperature for decontamination, and sometimes application of chemicals to prevent pest damage, which could further DNA degradation and damage (Brewer et al., 2019; Staats et al., 2011). The most common type of chemical damage in herbarium aDNA is cytosine deamination to uracil. While DNA in herbarium samples degrades 6–8 times faster than in archaeological bones of Moa birds, levels of cytosine deamination remain relatively low, reflecting its gradual, time-dependent nature (Allentoft et al., 2012; Kistler et al., 2017, 2020; Porrelli et al., 2026; Weiß et al., 2016). Additionally, DNA from herbarium specimens is typically a complex mixture from the targeted plants and metagenomic DNA of microorganisms present before death or introduced post-mortem, with the relative proportions of endogenous and exogenous DNA varying widely among samples (Gutaker & Burbano, 2017; Ng & Gutaker, 2025; Porrelli et al., 2026; Weiß et al., 2016).

Leaves have been the most commonly used tissue for DNA isolation from herbarium specimens, due to its ready availability and potentially high DNA content (Drábková, 2013; Särkinen et al., 2012). However, the quality and quantity of DNA isolated from herbarium leaves can vary widely among taxonomic groups and individual specimens, affected by factors including sample age, taxon, and the source herbarium (Brewer et al., 2019; Kates et al., 2021; Porrelli et al., 2026; Quatela et al., 2023). Additionally, leaf tissue is not always present in sufficient quantity, particularly for small plants or specimens with few leaves. These limitations have prompted the development of non-destructive sampling methods that minimise damage to irreplaceable material while still yielding enough endogenous DNA for downstream genomic analyses (Sugita et al., 2020).

Aside from leaves, many studies have investigated DNA isolation from non-viable archaeological and historical seeds, including rice and barley seeds or seed remains (Hagenblad et al., 2024; Hiraoka et al., 2009; Kobayashi et al., 2006; Leino et al., 2009; Lister et al., 2009; Mascher et al., 2016; Muto et al., 2020; Palmer et al., 2009). Additionally, a few investigations have attempted DNA isolation from aged or mummified seed embryos, using either single embryos or pools of up to 16 embryos from certain eudicot and gymnosperm species (Rogers & Bendich, 1985; Walters et al., 2006). DNA quality, assessed via PCR amplification or NGS, demonstrated that DNA could be successfully isolated from seeds dating back to the 7^th^ century. The extent of DNA degradation and contamination was influenced by both sample age and species, while the seed coat may provide additional protection against DNA damage (Gugerli et al., 2005; Leino et al., 2009; Walters et al., 2006). In a singular case of large seeds from date palm, DNA was intact sufficiently to reportedly allow germination from 2,000-year-old archaeological sample (Gros-Balthazard et al., 2021).

However, DNA isolation from whole seeds may also be hindered by surrounding storage compounds such as carbohydrates, lipids, and polyphenols, which can interfere with DNA isolation and reduce yield and purity (Liang et al., 2016). Additionally, a complicating factor is that most seeds contain tissues of different individuals and of various ploidy levels, with maternal seed coat (2n), mixed endosperm (3n) and embryo (2n). Only the embryo represents the true diploid genotype of the progeny and consist of densely packed cells with relatively lower storage and secondary metabolite compounds, making them a potentially better target for DNA isolation from historical plant materials (Bennett et al., 1975; Itoh et al., 2005).

This study assessed the feasibility of isolating and sequencing DNA from single seed embryos from historical plant samples for whole genome sequencing. To that end, we compare DNA quality between embryos and leaves from the 26 specimens from three species: *O. sativa*, *O. rufipogon*, and *H. spontaneum*, and evaluate metrics such as endogenous DNA content, median fragment length, damage fraction per site, decay rates, nucleotide misincorporations, and library complexity.

## MATERIALS AND METHODS

### Herbarium specimen sampling

Plant materials (leaves and seeds) were obtained from herbarium specimens stored in the Ethnobotanical collection of National Museum of Natural History, France (NMNH), and Royal Botanic Gardens, Kew, United Kingdom (RBGK). *Oryza sativa* (cultivated rice) and *O. rufipogon* (wild rice) samples had been collected in Vietnam and Thailand, respectively, while *Hordeum spontaneum* (wild barley) specimens had been collected in West Asia (Iraq, Jordan, Syria, and Turkey) and Afghanistan. Ages of all samples ranged approximately from 30 to 200 years old (**Supplementary Table 1**).

### DNA isolation, library preparation and sequencing

Rice and barley seeds obtained from herbarium specimens were de-husked, followed by embryo separation by Swann-Morton^TM^ sterile, single-use stainless steel surgical scalpels (Fisher Scientific, 12397999). Rice embryos were smaller and lighter than wild barley embryos (**Figure 1**).

**Figure 1.**
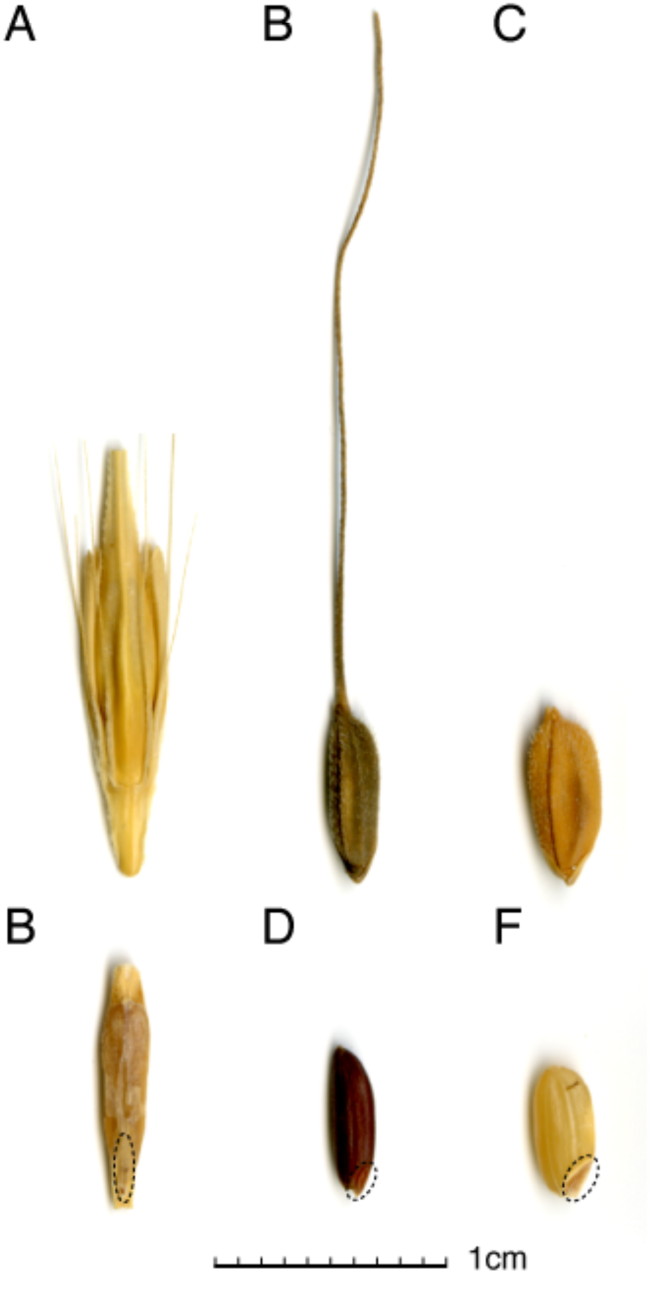
Representative husked and de-husked seeds of *H. spontaneum* (A, B), *O. rufipogon* (C, D) and *O. sativa* (E, F) sampled from herbarium specimens, dated to 1885, 1929, and 1914, respectively. Dashed lines encircle embryos in the de-husked seeds.

Dried tissues were subsampled into 2 mL PowerBead tubes (QIAGEN, 13117-50, metal 2.38 mm) and disrupted by using a Precellys Evolution homogeniser (2 – 4 cycles, soft program). Genomic DNA (gDNA) was isolated from dried leaf tissue (∼ 4 – 17 mg) and a single seed embryo using the previously published N-phenacylthiazolium bromide (PTB) – dithiothreitol (DTT) protocol (Gutaker et al., 2017; Latorre et al., 2020) in the clean room facilities at RBGK. In short, ground samples were lysed by resuspending in 1 mL of PTB lysis buffer (1% sodium lauryl sulphate (SDS), 10 mM Tris-HCl, 10 mM ethylenediaminetetraacetic acid (EDTA), 5 mM sodium chloride (NaCl), 2.5 mM PTB, 50 mM DTT, and 0.4 mg/mL Proteinase K) and incubating at 37°C for 16 – 24 hours in an incubated rotator. Lysate filtration and DNA purification were then carried out using a QIAGEN DNeasy Mini Plant kit (69106), following the manufacturer’s manual. gDNA was eluted from the DNeasy spin column in 100 µL buffer AE, and then quantified using a Quantus^TM^ Fluorometer system (Promega). A negative control was included in each batch of DNA isolation.

Sequencing libraries were prepared from 20 or 26 µL of purified gDNA (depending on gDNA concentrations, **Table 1**) using previously published protocols (Kircher et al., 2012; Latorre et al., 2020; Meyer & Kircher, 2010). Each batch of library preparation had a negative control (distilled water instead of DNA). The process consisted of four main steps (blunt-end repair, adapter ligation, adapter fill-in, and indexing PCR). Uracil-DNA glycosylase (UDG) treatment was not carried out during the blunt-end repair process; to maintain native levels of DNA damage patterns. Indexing oligo sequences were adapted from Meyer & Kircher (2010) and indexing PCR was run for 10-12 cycles. QIAGEN MinElute PCR Purification kit was employed to purify DNA after every step except for adapter fill-in where enzyme heat inactivation was performed. Indexed libraries were then eluted in 40 µL of Buffer EB, quantified and visualised by using Quantus^TM^ Fluorometer and 4200 TapeStation (D1000, Agilent) systems.

**Table 1.**
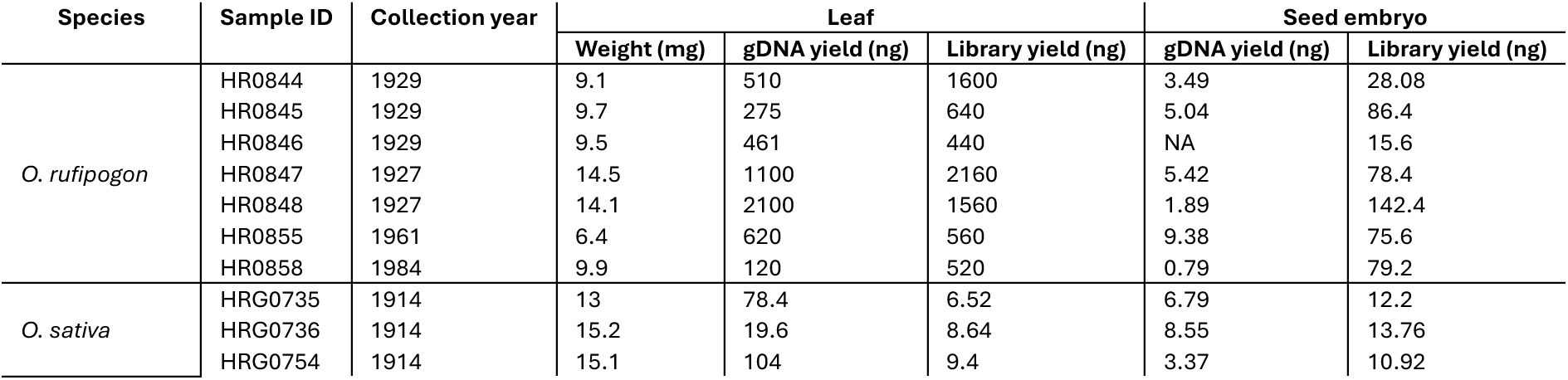
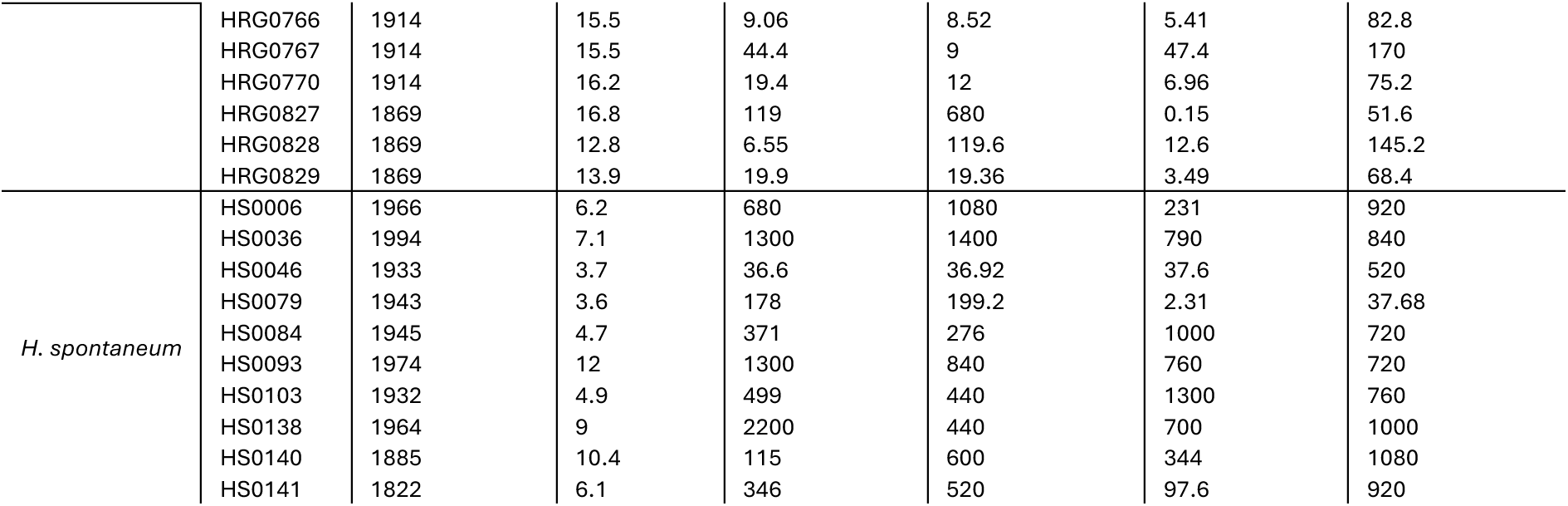
Yields of gDNA and sequencing libraries isolated from leaf or a single seed embryo sampled from *O. sativa*, *O. rufipogon*, and *H. spontaneum* herbarium specimens.

Generated libraries were then pooled and sequenced on an Illumina NovaSeq X+ system at Macrogen Europe B.V using chemistry for 150 cycles in paired-end mode.

### Bioinformatics analyses

Bioinformatic protocols for screening and determining properties of sequenced historical DNA libraries were adapted from Latorre et al. (2020) with some modifications, listed below.

#### Read processing and alignment, and library complexity

Raw demultiplexed paired reads were merged (minimum overlap of 11 base pairs) and adapter sequence was removed using AdapterRemoval v2.3.4 (Schubert et al., 2016). Quality check of trimmed and merged reads was then performed using FastQC v0.12.1 (*Babraham Bioinformatics - FastQC A Quality Control Tool for High Throughput Sequence Data*, n.d.). Subsequently, processed reads were mapped to their corresponding reference genomes using bwa-aln v0.7.18 (Li & Durbin, 2009) with disabled seed (option -l 1024). Reference sequences of *O. sativa, O. rufipogon*, and *H. spontaneum* were downloaded from National Center for Biotechnology Information (NCBI) (GCA_001433935.1, GCA_037997075.1, and GCA_949783245.1, respectively). Unmapped reads were then flagged and removed by using bwa-samse and samtools v1.21 (Danecek et al., 2021). Library complexity – the proportion of i) distinct reads to ii) mapped reads of each sequencing library was estimated by using preseq c-curve function (Daley & Smith, 2013). Due to low endogenous contents and low depth sequencing, most *O. sativa* leaf libraries had less than 20,000 reads post filtering (merging, quality check, and mapping), with the lowest library containing ∼4000 reads, while the majority of embryo libraries of three examined species as well as leaf libraries of *O. rufipogon* and *H. spontaneum* had at least 480,000 merged mapped reads, with the largest library comprising ∼53 million reads. C-curve complexity analysis showed unstable estimates at 4000 reads mark, and therefore, complexity output was reported at a fixed depth of 480,000 reads, when available. Finally, PCR optical duplicates were removed by using DeDup v0.12.9 (Peltzer et al., 2016) for downstream analysis.

#### DNA damage patterns and degradation analyses

##### Endogenous content

Endogenous DNA content of each sample was measured by the proportion of the number of mapped reads (primary alignments to its corresponding reference genome) to the total number of quality-control (QC)-passed reads. Samtools flagstat tool (Danecek et al., 2021) was used to extract this information from sam files. Endogenous contents across different tissue types within each species were then plotted using the ggplot2 package in R.

##### C-to-T misincorporations

Fractions of C-to-T misincorporations of the first 30 nucleotides (5’-end) were determined by using AMBER tool (Dolenz et al., 2024), and then visualised using the ggplot2 package in R.

##### Median fragment length

For each sample, read lengths and read counts per length were calculated by AMBER tool (Dolenz et al., 2024). As fragment length distributions were typically right-skewed, with many short fragments and fewer long fragments, a lognormal distribution was fitted to determine the median fragment length as well as standard deviation (SD) value for each sample. This analysis was carried out by using the fitdistr function (MASS package) in R following Weiß et al. (2016). Visualisation of these variations by tissue type was performed using ggplot2 package in R.

##### Damage fraction per site (λ) and decay rates (k)

The damage fraction per site (*λ*) of each sample was estimated by using a previously described method (Allentoft et al., 2012; Weiß et al., 2016). Briefly, the fragment length having the highest number of reads (peak fragment length) was determined, and all fragment lengths greater than or equal to this peak and up to 110 bp were isolated. This is to avoid the part of the distribution that is affected by the loss of small molecules during laboratory processing. A log-linear regression model was then fitted to this subset to determine the relationship between the fragment length and the logarithm value of the corresponding fragment frequency. The negative of this regression slope (*λ*) illustrated the fraction of DNA bonds per nucleotide in the sample – the rate at which the number of reads decreased with increasing fragment length. Linear regressions of *λ* versus sample age, grouped by tissue type and species, were then generated. Other statistical features including slope (*k*, DNA decay rate per base per year), *R^2^*, and *p*-value were also calculated to evaluate the relationship between DNA degradation and age in accordance with species and tissue characteristics.

## RESULTS

### DNA isolation and library preparation

Historical DNA was successfully isolated from leaves and individual seed embryos obtained from herbarium specimens of cultivated rice (*O. sativa*), wild rice (*O. rufipogon*), and wild barley (*H. spontaneum*), spanning approximately 30 to 200 years in age (**Table 1**). gDNA concentrations ranged from 0.0015 to 22 ng/µL, corresponding to total yields of 0.15 ng – 2.2 µg. No linear relationship was observed between the amount of leaf tissue used and the achieved gDNA yield (**Table 1**). Mass of embryos was below detection limit of laboratory scales and were not reported. Regardless of the significantly low gDNA yields of some samples (especially in embryo DNA samples), sufficient amounts of gDNA were isolated for downstream library preparation as well as whole genome sequencing (**Table 1**).

### Endogenous content, library complexity, nucleotide misincorporations, and DNA fragmentation

To estimate contamination levels and the proportion of endogenous DNA, processed DNA reads were mapped to their corresponding reference genomes. Seed embryos of all species (except for one wild barley sample), as well as leaves of *O. rufipogon* and *H. spontaneum* (except for two wild barley samples), showed high endogenous DNA content (83 – 98%). In contrast, endogenous percentages of *O. sativa* leaves were relatively low and variable, ranging from ∼1.6 – 37%. Accordingly, the endogenous DNA proportions were only significantly higher in *O. sativa* seed embryos compared to leaves (paired *t*-test, *p* = 4.5 x 10^-8^) (**Figure 2A**).

**Figure 2.**
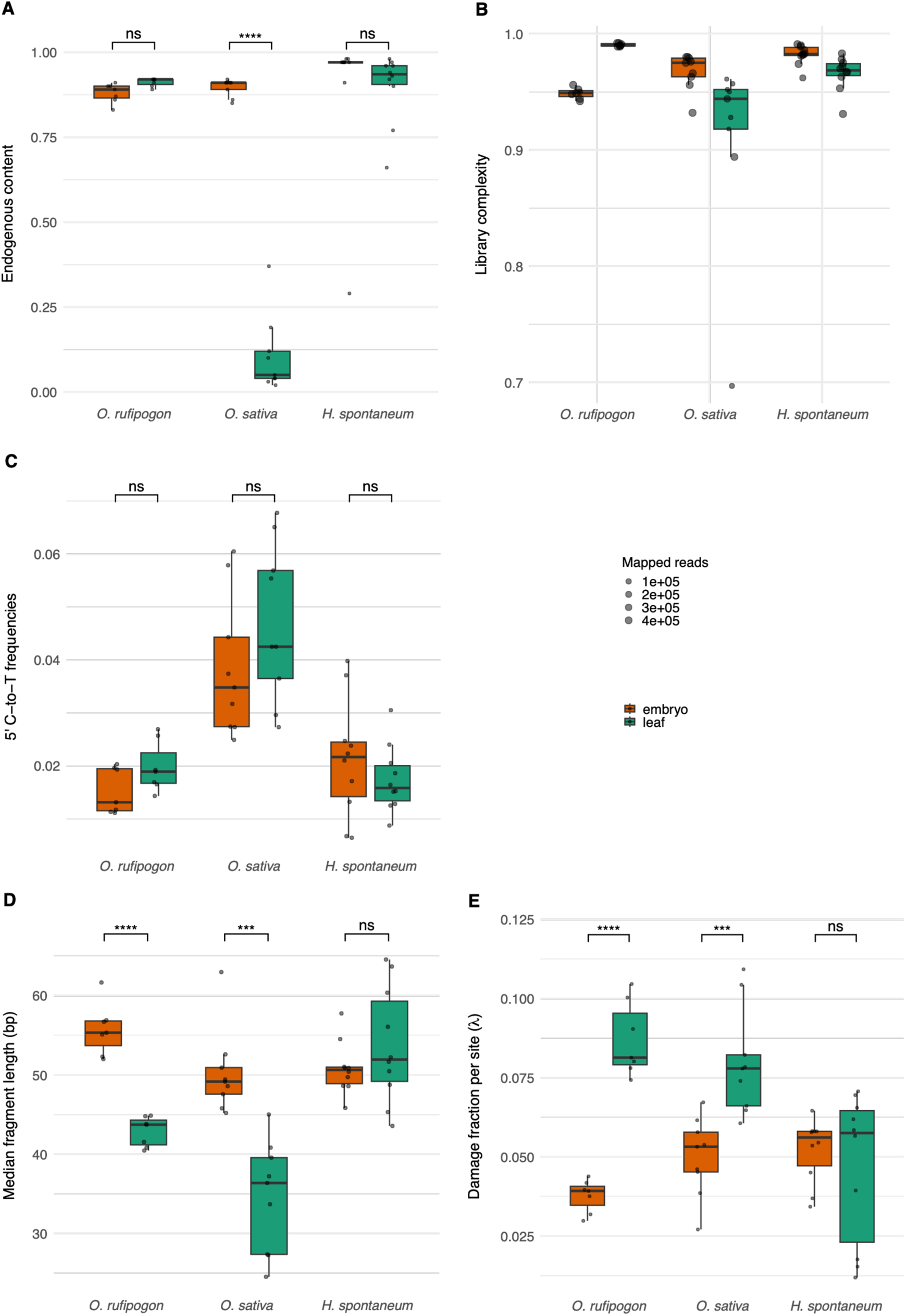
Effects of different tissue types on DNA preservation in historical samples. Box and whisker plots illustrate differences in endogenous content (A), library complexity (B), 5’ C-to-T frequencies at the first base of sequenced reads (C), median fragment length (D), and damage fraction per site (*λ*) (E) of DNA sequencing libraries of wild rice (*O. rufipogon*), cultivated rice (*O. sativa*), and wild barley (*H. spontaneum*). Paired *t*-tests were performed to compare DNA characteristic (A – D) in two tissue types (ns: not significant, ***: 0.0001 ≤ p ≤ 0.001, ****: p < 0.0001). Dot sizes in plot B are scaled by total number of merged mapped reads.

Sequenced library complexity was also assessed to validate the capability of very high-throughput sequencing typical for whole genome sequencing at high coverage for most downstream genomic analyses. Preseq extrapolation curves for both tissue types in *O. rufipogon* and *H. spontaneum* samples indicated that library complexities were consistently high, ranging from 93 to 99%. Cultivated rice *O. sativa* leaf libraries exhibited a wider complexity range (∼70 to 96%) (**Figure 2B**, **Supplementary Table 2**).

To evaluate spontaneous post-mortem modifications (DNA damage), the fraction of first-base C-to-T nucleotide misincorporation – the most common type of miscoding lesions (Briggs, 2007), were estimated. Overall, *O. sativa* samples exhibited higher C-to-T frequencies (up to 0.068) than most samples of the other two species (0.006 – 0.03). The differences in deamination rates between two tissue types were visible but statistically insignificant across all species (**Figure 2C**).

Median values, which were estimated from fitted lognormal distributions of merged reads, were used to investigate differences in DNA fragment lengths across tissues and species. Comparisons of these median lengths demonstrated a pronounced tissue-type effect in *Oryza* samples, with embryo samples showing substantially longer fragments than leaves in both *O. rufipogon* and *O. sativa* (paired *t*-test, *p* = 8.85 x 10^-5^ and *p* = 1.3 x 10^-4^, respectively). In contrast, no significant differences were observed between the two groups in wild barley (**Figure 2D**).

In conclusion, tissue type had a significant impact on both the quantity and quality of retrieved endogenous DNA from herbarium specimens of examined *Oryza* species, although in *O. rufipogon* only median DNA fragment length was improved in embryo vs. leaf. On the other hand, satisfactory yields and quality of endogenous DNA for whole genome sequencing were isolated efficiently from *H. spontaneum* in both tissue types.

Fractions of intact bonds per nucleotide (*λ*) were calculated for all samples, followed by paired *t*-tests between two tissue types. The results revealed that leaves in both *O. rufipogon* and *O. sativa* exhibited significant higher damage fractions per site than embryos (*p* = 2.38 x 10^-5^ and 6.38 x 10^-4^, respectively), whereas insignificant differences in *λ* values were found between two tissue types in *H. spontaneum* (**Figure 2E**).

DNA decay rates per base per year (*k*) were then estimated by plotting λ value against sample age. Across all three species, *k* values of DNA retrieved from leaf tissue were consistently higher than those from corresponding embryo tissue, indicating faster decay rates in leaves, although in many cases the regression correlations between λ and age were weak or not statistically significant. In *O. rufipogon*, embryos showed a strong and statistically significant supported age – damage relationship (*R^2^* = 0.825, *p* = 0.005), whereas leaves displayed a weaker correlation (**Figure 3A**). In *O. sativa*, leaf tissue exhibited both a higher decay rate and a statistically significant positive correlation with age (*k* = 5.86 x 10^-4^, *R^2^* = 0.606, *p* = 0.013), while seed embryo tissue had a lower and less consistent decay pattern (**Figure 3B**). In *H. spontaneum*, decay rates in both tissues were relatively lower (embryo: *k* = 1.05 x 10^-4^ and leaf: *k* = 2.58 x 10^-4^) than those of *Oryza* species, and *λ*-age correlations were weak and not significant, suggesting minimal influence of tissue type on DNA degradation in this species (**Figure 3C**).

**Figure 3.**
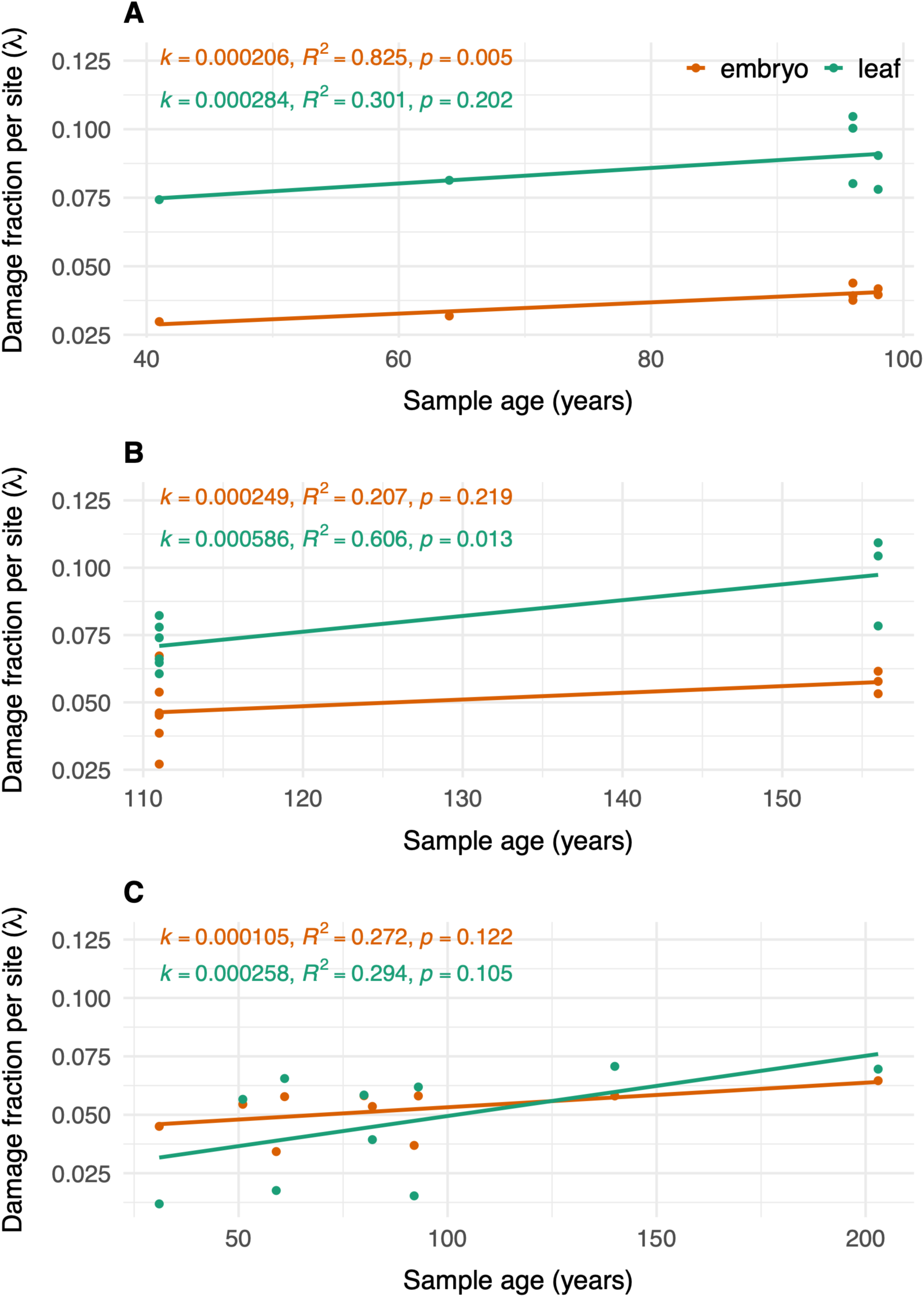
Relationship between damage fraction per site (*λ*) and sample age of DNA. Scatter plots for *λ* values and age in two tissue types across the three investigated species *O. rufipogon* (A), *O. sativa* (B), and *H. spontaneum* (C).

Accordingly, these results demonstrated that tissue type substantially affected DNA decay rates in investigated *Oryza* samples, with leaves generally decaying faster than embryos, whereas this effect was not significant in *H. spontaneum*.

### Assessment of historical crop seed materials in global collections

Herbarium sheets might have seeds attached to the specimens, in special capsules or in associated carpological collection. However, seeds are not always present - botanists prefer flowering stage for most species, which often preserves more taxonomy-informative characteristics. By contrast, biocultural collections usually focus on the utilised portion of the plant, which is the seed or fruit in the case of many crop plants. Many biocultural collections were formed in the eighteenth and nineteenth centuries as research resources in the fields of economic botany and ethnobotany, agronomy, pharmacognosy, food science and others related to useful plants.

Herbarium materials for genetic analysis can be identified either through Index *Herbariorum*, or via the global compendium of specimen records at the *Global Bioinformatics Information Facility* (GBIF). By contrast, biocultural collections are usually poorly known and incompletely catalogued, even when stored at the same institutions as herbaria (Cornish & Nesbitt, 2014; Nesbitt & Cornish, 2016). The Biocultural Collections Group, a specialist network within the Society for Ethnobotany, has done much to bring curators together, but a comprehensive census of such collections is still far from completion (Salick et al., 2014). We draw on our networks to highlight a representative sample of 25 biocultural collections, indicative of those available worldwide. Three largest collections with samples from global distribution are at the Natural History Museum in Paris, the Royal Botanic Gardens, Kew, and Vavilov Institute of Plant Industry in Saint Petersburg (**Table 3**). Most smaller collections offer research resources, more focused on particular crops or geographies (**Table 3**). Samples in biocultural collections can be housed in herbarium format, attached to paper mounts but emphasising reproductive parts. Alternatively, collections are housed in glass jars, plastic boxes and similar containers (**Figure 4**).

**Figure 4.**
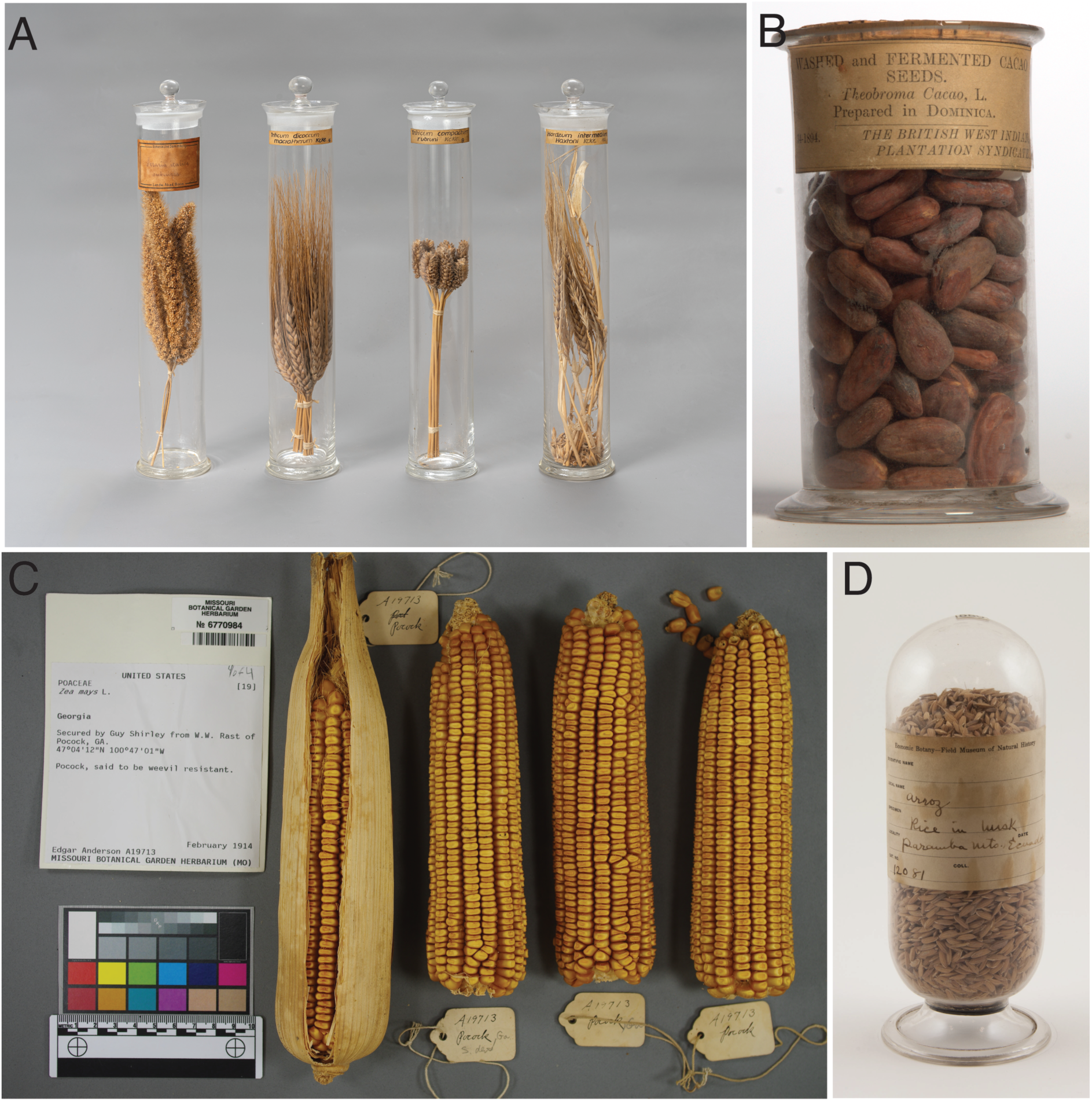
Examples of biocultural collection specimens with crop seeds. Ears of foxtail millet (*Setaria italica*), emmer wheat (*Triticum dicoccum*), bread wheat (*Triticum compactum*); barley (*Hordeum intermedium*), c. 1870, Körnicke collection, University of Bonn (A); CC BY 4.0. Cacao beans, 1894, Dominica, Royal Botanic Gardens, Kew (EBC 070828) (B); CC BY. Yellow dent corn (*Zea mays*) from Georgia, USA, 1914, Missouri Botanic Garden (MO-3056378) (C); CC BY-NC-SA 3.0. Rice (*Oryza sativa*) grains, Ecuador, Field Museum (Economic Botany 12081) (D); CC BY-NC 4.0

**Table 3.** Representative table of biocultural collections known to have significant holdings of crop seeds.

| Collection | Approx. number crops | Date | Specialisms | Country |
| --- | --- | --- | --- | --- |
| Powerhouse Museum, Sydney | 100+ | 1850s- | Global, Australia | Australia |
| Economic Botany Collection<br>Meise Botanic Garden | 1000+ | 1821- | Global, Africa | Belgium |
| Canada Agriculture and Food Museum, Ottawa | 100+ | 1800-2000 | Canada | Canada |
| Biocultural Collection Natural History Museum, Copenhagen | 500+ | 1850-1950 | Global | Denmark |
| Ethnobotany collection, National Museum of Natural History, Paris | 80000+ | 1911-1970s | Southeast Asia and global. Rice, coffee, sorghum, millets. | France |
| Ethnoecology collection, National Museum of Natural History, Paris | 100+ | 1960s- | Global | France |
| Botanische Gärten der Universität Bonn | 1000+ | 1867-1908 | Global. Cereals collection of Friedrich August Körnicke. | Germany |
| Applied Botany Collection (ABC), Biozentrum Klein Flottbek, University of Hamburg | 1000+ | 1870-1970 | Global | Germany |
| Archaeobotany Laboratory, Bar-Ilan University | 1000+ | 1870s- | Israel and Palestine. Includes wheat landrace collection of Dr. Ludwig Pinner. | Israel |
| Herbarium, Museo di storia naturale dell'Università degli studi di Firenze | 100+ | 1912-1924 | Global, Africa | Italy |
| Museo agrario tropicale dell'istituto agronomico per Oltremare, Florence | 100+ | 1904-1970s | Africa | Italy |
| Hideo Hamada collection, Kobe University | 1000+ | 1957-1958 | Southeast Asia. Rice. | Japan |
| Ethnobotany Collection, Institute of Biology, UNAM | 100+ | 1980s- | Mexico | Mexico |
| Economic Botany Collection, Naturalis, Leiden | 1000+ | 1800- | Global | Netherlands |
| Colonial Agricultural Museum, Natural History Museum, Lisbon | 100+ | 1850- | Africa | Portugal |
| Economic Botany Collection, Komarov Botanical Institute, St Petersburg | 1000+ | 1800- | Global | Russia |
| Seed collection, Vavilov Institute of Plant Industry, St Petersburg | 10000+ | 1890- | Global. Distinct from the seedbank collection. | Russia |
| Botanical Collection, Natural History Museum, University of Zürich | 1000+ | 1850s- | Global | Switzerland |
| Herbarium, Royal Botanic Gardens, Kew. | 1000+ | 20<sup>th</sup> century | Africa. Snowden collection of sorghum. Also rice, millet etc. These specialist collections with emphasis on crop landraces are held separate to main Herbarium. | United Kingdom |
| Economic Botany Collection, Royal Botanic Gardens, Kew | 10000+ | 1847- | Global | United Kingdom |
| John Percival Collection, Herbarium, University of Reading | 5000+ | <1921 | Eurasia. Wheat. Smaller sets are held by Kew, the Natural History Museum, London and the Vavilov Institute, St Petersburg. | United Kingdom |
| Timothy C. Plowman Economic Botany Collection, Field Museum, Chicago | 1000+ | 1893- | Global, Americas | United States |
| Biocultural Collection, Missouri Botanical Garden | 1000+ | 1900- | Americas. Maize. | United States |
| Economic Botany Collection, Harvard University Herbaria | 1000+ | 1858-1970s | Global, Americas | United States |
| Biocultural Collection, New York Botanical Garden | 1000+ | 1891-1970s | Global, Americas | United States |

## DISCUSSION

The main aim of this study was to test the possibility of whole-genome sequencing from plant embryos and compare the aDNA characteristics of sequenced libraries generated from embryos to those of leaf tissue across three species, *O. rufipogon*, *O. sativa*, and *H. spontaneum*. The results have demonstrated the successful recovery of gDNA from both tissue types, and remarkably, from a single seed embryo of up to ∼200 years old specimen. Although gDNA yields were low in some samples, libraries were generated and sequenced efficiently with DNA mixture complexity that would easily allow sequencing whole genomes to a desired depth of coverage. While several previous studies using herbarium materials reported that the DNA inputs of 500 pg to ∼ 10 ng were adequate for genome skimming and targeted enrichment sequencing (Hart et al., 2016; Zeng et al., 2018), sequencing libraries in this study were constructed from as little as ∼40 pg of degraded aDNA, providing sufficient quantities for whole-genome sequencing.

To further evaluate the quality of retrieved aDNA, endogenous content, library complexity, fragment length, and damage profiles were estimated. Several previous studies have shown that endogenous DNA proportion was primarily influenced by environmental conditions, including annual precipitation and temperature seasonality, rather than by sample age (Brewer et al., 2019). Differences in tissue properties (e.g., tissue texture, secondary metabolites) approximated by diverse taxonomic groups, also impacted the aDNA preservation in herbarium specimens (Brewer et al., 2019; Kates et al., 2021). Finally, endogenous DNA can be rapidly depleted due to cellular processes and microbial digestion, as well as specimen preparation (for example oven-drying). Among the three examined species in this study, both *Oryza* species were collected in a wet tropical climate and had similar tissue characteristics. Despite this, we observed very low endogenous DNA content only in *O. sativa* leaf samples, indicating that i) idiosyncratic processes, such as specimen preparation differentiated it from *O. rufipogon* leaf samples, and ii) such processes reduced DNA quality in leaf tissue, but not in embryo. This suggested that herbarium preparation process in *O. sativa* likely caused substantial DNA loss in leaves, whereas the relatively thick seed husk provided structural protection (Gugerli et al., 2005), preserving endogenous DNA in the embryo. However, the lack of detailed preparation and preservation histories for most herbarium specimens has limited the ability to draw strong conclusions about the impact of processing methods on DNA integrity (Brewer et al., 2019; Staats et al., 2011).

In addition, tissue type effects were observed in both *Oryza* species with respect to the median DNA fragment length. Rapid DNA degradation and fragmentation following organismal death are primarily driven by intracellular nucleases and microbial activities, and these processes are tissue dependent. Subsequently, DNA fragmentation proceeds at a slower, more constant rate, likely caused by depurination (Briggs et al., 2007; Lindahl, 1993) and scales with passing time (Porrelli et al., 2026; Weiß et al., 2016). DNA fragmentation is only minimally affected by the climate during collection (Porrelli et al., 2026). The protective effect of the seed coat could have mitigated DNA degradation, resulting in longer median fragment lengths in embryo-derived sequencing data compared with leaf tissue in tropical *O. sativa* and *O. rufipogon*. Additionally, rice seeds tolerate drying (Choudhary et al., 2023) – the crucial step in herbarium specimen preparation (Bridson & Forman, 1998) – and can thus retain seed embryo viability longer than vegetative tissues such as leaves. Consequently, DNA fragmentation and damage may accumulate more slowly in seed embryos than in leaves from the same plant. Conversely, *H. spontaneum* samples collected from temperate environments may have experienced less severe DNA fragmentation in leaf tissue and hence the difference in median length of sequenced DNA compared to embryo is non-significant. In line with this, the damage fraction per site (*λ*) also differed between tissue types in *Oryza* species but not in *H. spontaneum*. The findings of this study further suggest that the DNA fragmentation in herbarium samples collected from tropics could be partly mitigated by the protective properties of the seed husk.

Our study opens up new avenues for research on historical plant materials, particularly crops. Herbaria are hugely promising sources of study materials for plant evolution and ecology, however, herbarium specimens have limitations: the amount of plant material available for sampling may be limited, crops are typically under-collected by botanists, and specimens often do not bear cultural data such as the name of the landrace or cultivar. Conversely, specimens in biocultural collections offer significant advantages to genetics researchers. The emphasis is on the utilised part of the plant, thus seeds and fruits in the case of many crops (**Figure 4**). They are often in bulk, enabling easier decision making on destructive sampling, and often bear significant cultural data.

These specimens often benefit from the significant investment in researching provenance and connections to previous publications, which will require advice from curators or historians of science. Compared to genebank collections, seeds in biocultural collections are usually no longer viable, but have typically been stored in conditions that preserve aDNA in reasonable condition. Increased interest in sustainability and plant-human relations has led to a revival of interest in historic collections, and the formation of new ones (Salick et al., 2014). The approach reported in this study will facilitate the utilisation of genetic data from these rich and unique repositories.

Researchers will need to be creative in searching for such resources. They should bear in mind that the scope of historic collections is often wider than expected: for example, pharmacists usually had to study a wide range of food, dye and textile plants so these are well-represented in *materia medica* collections. Art or historical museums may hold substantial biocultural collections deriving from past World’s Fairs. We recommend that researchers begin with local collections and make wide enquiries. Use of such collections is key to increased societal value and, thus, increased investment in their curation. Further advice on their use is given by (Allasi Canales et al., 2022).

In conclusion, this study has demonstrated the feasibility of isolating DNA and constructing sequencing libraries from single seed embryos of *O. sativa*, *O. rufipogon* and *H. spontaneum*. In *Oryza* genus, embryos consistently outperformed leaves across several measures, whereas in wild barley both tissues delivered satisfactory results without significant differences. Embryos represent a promising alternative source for DNA isolation from herbarium specimens, where collection, preparation, and processing possibly severely compromise endogenous DNA quality. These findings highlight the broader potential for applying single-seed embryo DNA isolation in herbaria, particularly when leaf tissue is limited. Additionally, this research opens up previously underutilised sources of rich historical collections such as seed banks, economic botany collections, anthropological museums, as well as other collections where crop grains are preserved, for genomic investigations.

