## Supplementary material for "Historical plant embryos as alternative sources of ancient DNA for whole genome sequencing": Supp. Table 1

**Supplementary Table 1**. Details of herbarium samples (NMNH: National Museum of Natural History, France; RBGK: Royal Botanic Gardens, Kew, United Kingdom; TBU: To Be Updated).

| **Species** | **Sample ID** | **Herbarium** | **Herbarium ID** | **Collector** | **Collection year** | **Country** | **Sampling location** |
| --- | --- | --- | --- | --- | --- | --- | --- |
| *O. sativa* | HRG0735 | NMNH | TBU | P. Morange | 1914 | Vietnam | Ben Tre |
|  | HRG0736 |  | TBU | P. Morange | 1914 |  | Ben Tre |
|  | HRG0754 |  | TBU | P. Morange | 1914 |  | Ben Tre |
|  | HRG0766 |  | TBU | P. Morange | 1914 |  | Long An |
|  | HRG0767 |  | TBU | P. Morange | 1914 |  | Long An |
|  | HRG0770 |  | TBU | P. Morange | 1914 |  | Long An |
|  | HRG0827 |  | P01934491 | J. B. L. Pierre | 1869 |  | Tien Giang |
|  | HRG0828 |  | P01934492 | J. B. L. Pierre | 1869 |  | Tien Giang |
|  | HRG0829 |  | P01829496 | J. B. L. Pierre | 1869 |  | Tien Giang |
| *O. rufipogon* | HRG0844 | RBGK | K000631586 | N. Put | 1929 | Thailand | Angtawng |
|  | HRG0845 |  | K000631587 | N. Put | 1929 |  | Angtawng |
|  | HRG0846 |  | K000631588 | N. Put | 1929 |  | Angtawng |
|  | HRG0847 |  | K000631589 | D. J. Collins | 1927 |  | Lemchabang, Sriracha |
|  | HRG0848 |  | K000631590 | D. J. Collins | 1927 |  | Lemchabang, Sriracha |
|  | HRG0855 |  | K000682212 | Kai Larsen | 1961 |  | Ban Kao |
|  | HRG0858 |  | K000682218 | H. D. Catling | 1984 |  | Prajuburi |
| *H. spontaneum* | HS0006 | RBGK | K001451889 | Davis (42355) | 1966 | Turkey | Urfa |
|  | HS0036 |  | K001451941 | R.M.A. Nesbitt | 1994 | Israel | Giv’at Yo’av |
|  | HS0046 |  | K001451924 | F.S. Meyers & J. D. Dinsmore | 1933 | Syria | Daraa |
|  | HS0079 |  | K001451947 | P.H. Davis (6113 AA) | 1943 | Syria | NA |
|  | HS0084 |  | K001451952 | Davis (9378 B) | 1945 | Jordan | Zerka (Zarqa) |
|  | HS0093 |  | K001451959 | L.J.G. van der Maesen | 1974 | Iraq | Ninawa (Nineveh) |
|  | HS0103 |  | K001451969 | Meleki Beg | 1932 | Iraq | Duhok (Dohuk) |
|  | HS0138 |  | K001453021 | P. Furse | 1964 | Afghanistan | Kataghan (Qataghan) |
|  | HS0140 |  | K001453019 | J.E.T. Aitchison | 1885 | Afghanistan | Badghis |
|  | HS0141 |  | K001453017 | NA | 1822 | Afghanistan | NA |
